# A new class of constitutively active super-enhancers is associated with fast recovery of 3D chromatin loops

**DOI:** 10.1101/458117

**Authors:** Jayoung Ryu, Hyunwoong Kim, Dongchan Yang, Andrew J. Lee, Inkyung Jung

**Author notes:** These authors contributed equally to this work (Jayoung Ryu, Hyunwoong Kim). Correspondence to Inkyung Jung Email Addresses: JR, HK, DY, AJL, IJ.

## Abstract

Super-enhancers or stretch enhancers are clusters of active enhancers that often coordinate cell-type specific gene regulation. However, little is known about the function of super-enhancers beyond gene regulation. In this study, through a comprehensive analysis of super-enhancers in 30 human cell/tissue types, we identified a new class of super-enhancers which are constitutively active across most cell/tissue types. These ‘common’ super-enhancers are associated with universally highly expressed genes in contrast to the canonical definition of super-enhancers that assert cell-type specific gene regulation. In addition, the genome sequence of these super-enhancers is highly conserved by evolution and among humans, advocating their universal function in genome regulation. Integrative analysis of 3D chromatin loops demonstrates that, in comparison to the cell-type specific super-enhancers, the cell-type common super-enhancers present a striking association with rapidly recovering loops. We propose that a new class of super-enhancers may play an important role in the early establishment of 3D chromatin structure.

**Background:** Super-enhancers or stretch enhancers are defined by a strong enrichment of mediators and transcription-regulating proteins, appearing to play a deterministic role in cellular identity by controlling the expression of cell-type specific genes[1, 2]. Previous studies have revealed the critical function of super-enhancers during development and differentiation[3]. The enrichment of disease-associated single nucleotide polymorphism (SNP) in super-enhancers compared to that of typical enhancers proposed a substantial link between super-enhancers and many complex human diseases[2]. In addition, a set of recent studies have proposed potential functions of super-enhancers in the extremely long-range chromatin communications and the establishment of 3D chromatin loops[4]. These results suggest a more universal role of super-enhancers in genome regulation apart from cell-type specific gene regulation, but little is known about the mechanisms underlying these various functions. To extend the current knowledge of super-enhancers and their biological roles, we conducted a comprehensive analysis of super-enhancer activities across 30 human cell/tissue types. Our analysis suggests that a substantial number of super-enhancers exhibits prevalent activities across cell-types in terms of H3K27ac signals, and that these non-canonical super-enhancers are involved in the formation of fast recovering chromatin loops.

**Note:** A genome browser session has been set up for visualization of the super-enhancer domains described in the current study– https://genome.ucsc.edu/cgi-bin/hgTracks?hgS-doOtherUser=submit&hgS-otherUserName=abundantiavosliberabit&hgS-otherUserSessionName=SuperEnhancerDomain

## Results

### Genomic landscape of super-enhancer domains

To characterize different modes of super-enhancers across various human cell/tissue types, we first investigated the genomic distribution of super-enhancers across 30 human cell/tissue types, including 5 H1-derived early lineages, 8 immortalized cell lines, and 17 human postmortem tissues. Super-enhancers were defined using ROSE algorithm[2, 5] based on H3K27ac ChIP-seq signal (see Methods), which resulted in 17,916 super-enhancers with an average of 597 super-enhancers per cell/tissue type. Merging the super-enhancers in all of the cell/tissue types resulted in 6,039 putative super-enhancer regions, named super-enhancer domains (SE domains) (Figure 1a, see Methods). The super-enhancer domains presented a mean length of 32Kb, which was slightly longer than the individual super-enhancers with the mean length of 26Kb. Excluding any one of 30 samples did not significantly affect the overall distribution of length of super-enhancer domains distribution (one-way Analysis of Variance, p-value > 0.5), which excludes the possibility of a critical bias in the definition of super-enhancer domains caused by one or more of samples. In total, around 6.32% of the human genome was marked by super-enhancer domains, and 48.3% of the super-enhancer domains consisted of multiple super-enhancers identified in at least two or more cell/tissue types analyzed, suggesting a recurrent formation of super-enhancers in specific genomic regions.

**Figure 1.**
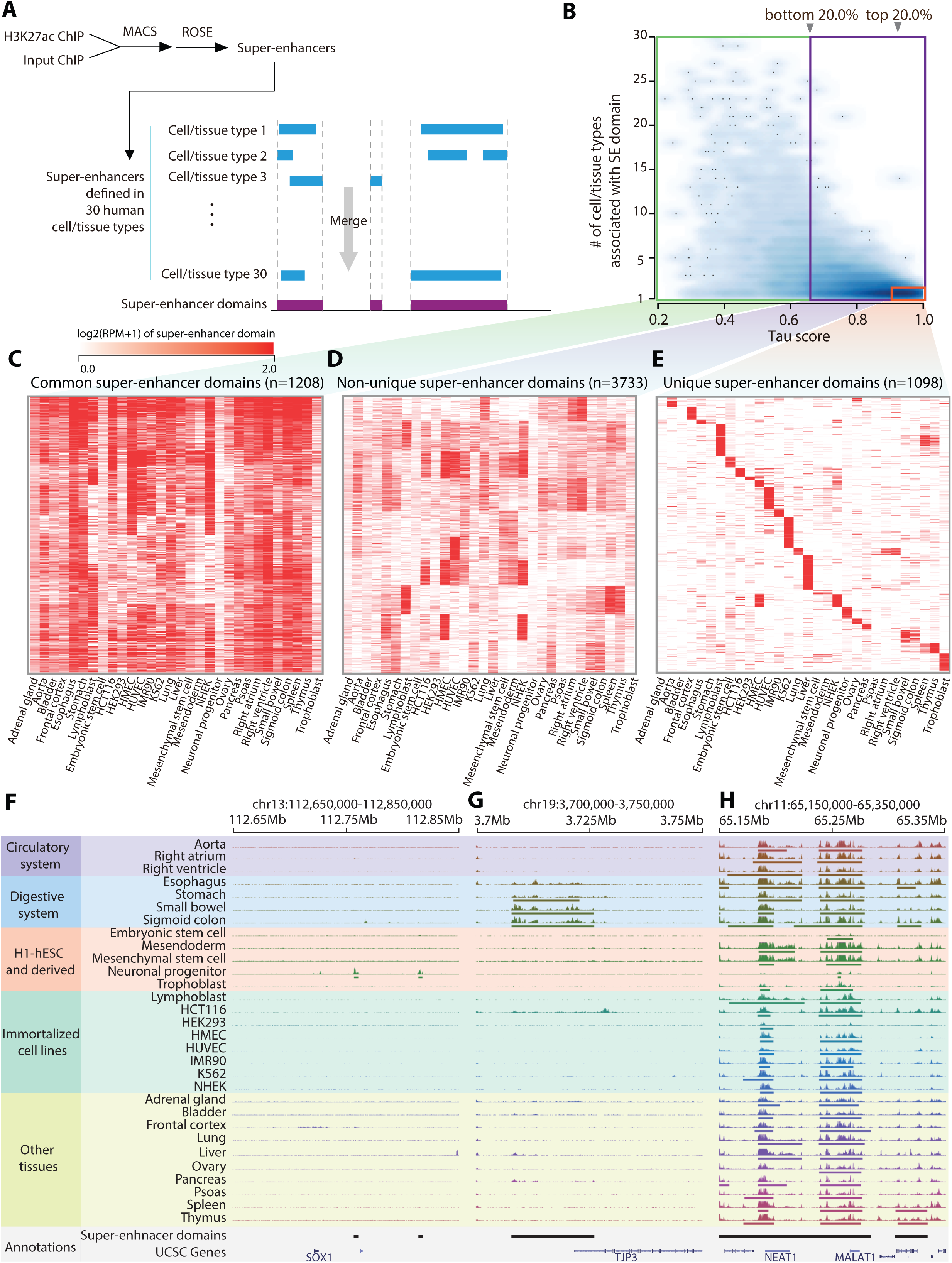
Super-enhancer domains can be grouped into distinct categories. **a,** A definition of super-enhancer domains. **b,** Smooth scatter plot of tau score (x-axis) and the number of cell/tissue types associated with super-enhancer domains. Vertical arrowheads indicate 20% and 80% percentile of tau score. Red, blue, and green colored box indicate unique, non-unique and common super-enhancer domains, respectively. **c-e,** Heatmaps of background-subtracted H3K27ac RPM of super-enhancer domains across cell/tissue types. Color intensity indicates log2 transformed H3K27ac RPM of super-enhancer domains. **f-h,** Genome browser snapshots showing H3K27ac signals of each class of super-enhancer domain examples. Unique super-enhancer domain nearby SOX1, specific to neuronal progenitor cells (f). Non-unique super-enhancer domain nearby TJP3, specific to digestive system related tissues (g). Common super-enhancer domain nearby MALAT and NEAT1 (h). H3K27ac signal is uniformly scaled in the range of 0-20. Cell/tissue types are grouped according to their lineage or functional similarity.

### Identification of common super-enhancer domains

In contrast to the previous notion that super-enhancers are highly cell/tissue-type specific, some super-enhancer domains exhibited a strong enrichment of H3K27ac signals in a surprisingly large number of cell/tissue types. To assess cell/tissue-type specificity of the super-enhancer domains, we utilized tau score[6], a measurement of tissue-specificity commonly used in gene expression studies (see Methods). Tau score of 0 indicates ubiquitous super-enhancer domain activity, while 1 represents tissue specific activation of the domain. The super-enhancer domain activities based on H3K27ac signals showed a wide spectrum of tau scores, reflecting a varying degree of cell/tissue-type specificity for each super-enhancer domain (Figure 1b). It should be noted that some super-enhancer domains were annotated by only one cell/tissue type, but showed low tau scores. Through manual inspection, we found that these domains presented moderately high H3K27ac signals in other cell/tissue types (Supplementary Figure 1), but the signals were insufficient to pass the threshold for super-enhancer calling by a small margin. For this reason, tau score was used as the main factor to define super-enhancer specificity.

In order to categorize super-enhancer domains with different cell/tissue-type specificity, we defined cell/tissue-type common and cell/tissue-type unique super-enhancer domains. The domains located within 20% or lower percentile of tau score were defined as common super-enhancer domains (1,208 domains, 20.0%, Figure 1c). On the other hand, domains that were called only in one cell/tissue type and located at 80% or higher percentile of tau score were defined as unique super-enhancer domains (1,098 domains, 18.2%, Figure 1e). Remaining domains were defined to be non-unique super-enhancer domains (3,733 domains, 61.8%, Figure 1d). To ensure that cancer cell lines or other immortalized cell lines did not produce critical bias by producing a large portion of common super-enhancers, we conducted hierarchical clustering between samples using the H3K27ac signals in the super-enhancer domains of available cell/tissue types (Supplementary Figure 2). The result demonstrated that, while tissue samples with close relatedness grouped together, cancer cell lines or other immortalized cell lines were not concentrated in the same cluster.

A notable example of unique super-enhancer domains was found in neural progenitor cells derived from human embryonic stem cells (H1-ESC). Two super-enhancer domains were uniquely defined in neural progenitor cells at downstream of SOX1 (Figure 1f), a gene known to be highly expressed only in neural progenitor cells and to play an important role in nervous system development. Non-unique super-enhancer domains showed high activity in several cell/tissue types but low activity in the rest. For example, a non-unique super-enhancer domain was called for tissues associated with the digestive system such as stomach, small bowel, and sigmoid colon (Figure 1g). There was also a high H3K27ac signal in the same domain in esophagus tissue, although it was not called as a super-enhancer. The putative target gene of the domain is TJP3 (tight junction protein 3), which is involved in junctional integrity of the intestinal cells. This example suggests a possibility that the formation of non-unique super-enhancer domains is a key regulator for shared functions in several tissues. Common super-enhancer domains showed high ubiquitous H3K27ac signals across the cell/tissue types, discordant with the previous notion of cell/tissue-type specificity of super-enhancers. An example of common super-enhancer domains was found near the noncoding RNA genes, NEAT1 and MALAT1, which are highly expressed across all cell/tissue types and previously suggested to function in a general biological process, such as the association with nuclear speckle formation[7] (Figure 1h).

### Distinct biological function of target genes in common super-enhancer domains

Super-enhancers are presumably associated with cell/tissue-type specific gene regulation, but the identification of distinct classes of super-enhancer domains raises the possibility that distinct modes of action exist for super-enhancers. We investigated the expression patterns of putative target genes in each class of super-enhancer domains. As expected, putative target genes of unique super-enhancer domains were highly expressed in the corresponding cell/tissue type (Figure 2a and b, see Methods). In contrast, genes associated with common super-enhancer domains showed universally high expression (Figure 2a and b). Although there was a significant enrichment of housekeeping genes[8] in putative target genes of common super-enhancer domains, the majority (82.4%) were non-housekeeping genes, indicating that housekeeping genes cannot fully explain the function of common super-enhancer domains. Gene set enrichment analysis of genes associated with unique super-enhancer domains showed enrichment in cellular identity functions in the corresponding cell/tissue type (Table 1, see Methods), whereas genes associated with common super-enhancer domains were enriched in basic cellular functions, such as transcriptional regulation, cell motility, and regulation of cell proliferation (Table 2). The overall pattern of expression and the significance of enriched pathways were moderately affected, but largely stayed consistent, when an alternative definition of putative target genes was used including the nearest 3 genes from the midpoint of each super-enhancer domain (Supplementary Figure 3a and b, Supplementary Table 1, 2). Our analysis revealed distinct biological functional enrichment in putative target genes between common and unique super-enhancer domains.

**Figure 2.**
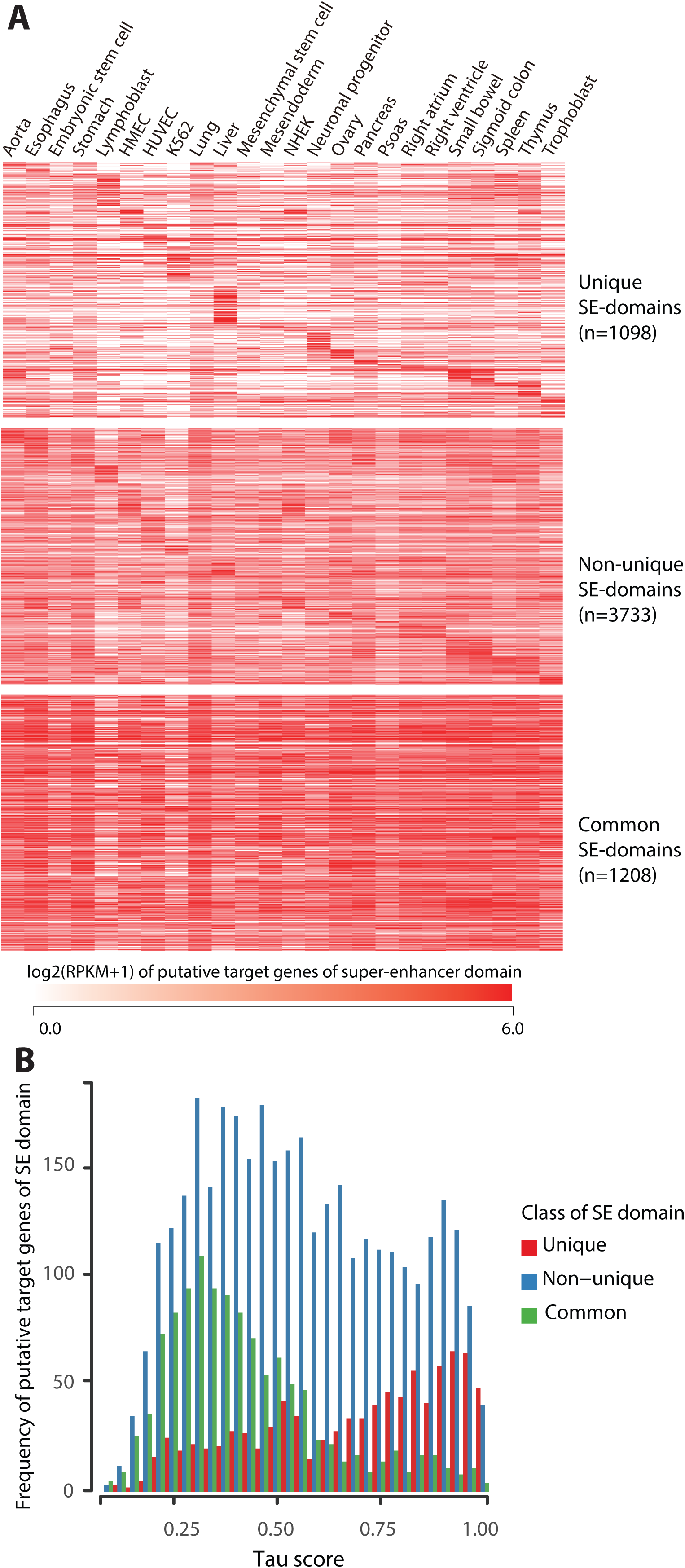
Functional characterization of super-enhancer domain classes. **a,** Heatmaps showing log2(RPKM+1) value of putative target gene of unique super-enhancer domains (top), non-unique super-enhancer domains (middle), and common super-enhancer domains (bottom). **b,** Histogram of tau score for putative target gene expression in each class of super-enhancer domains. Low tau score indicates universal expression pattern.

**Table 1.**
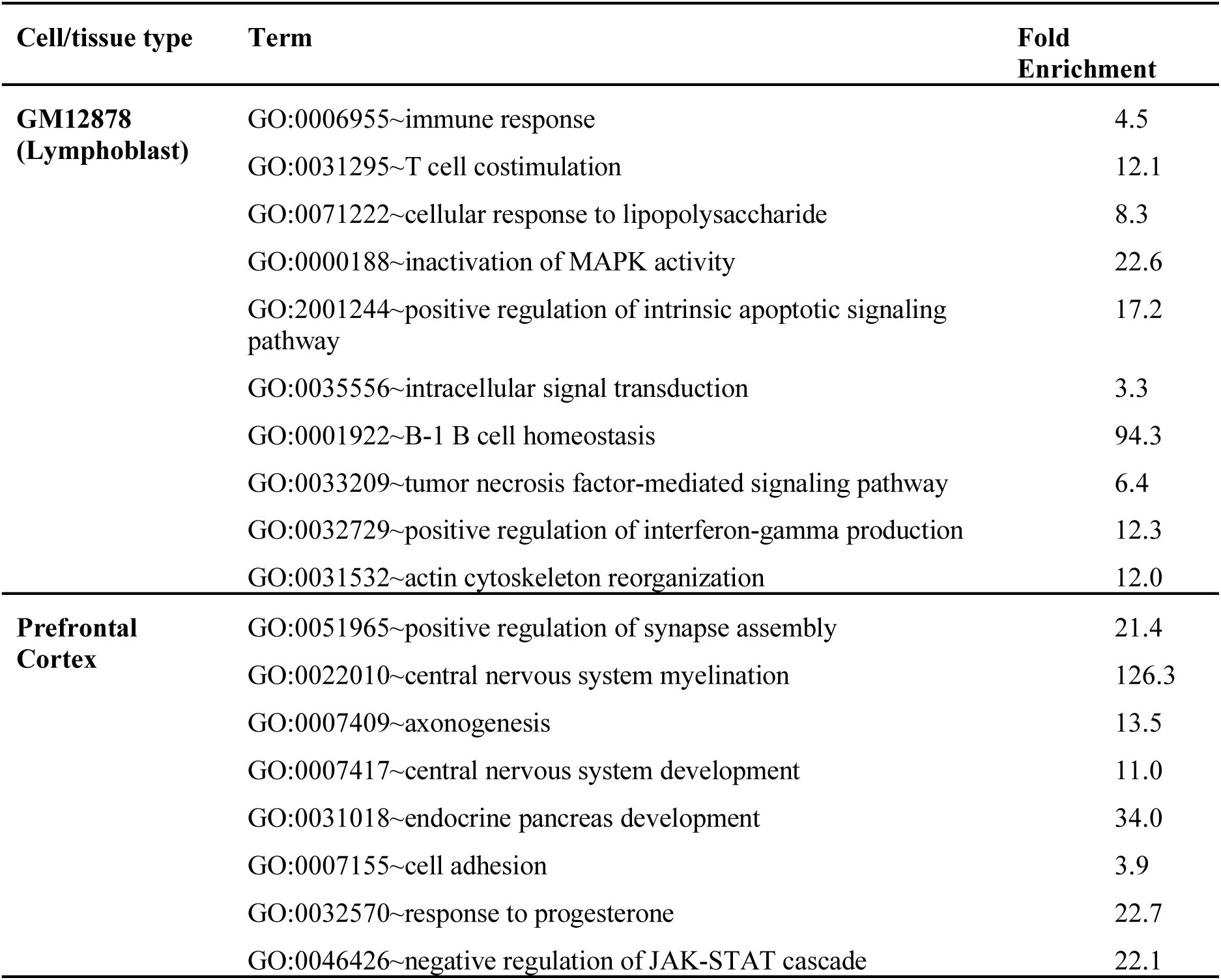
GO analysis of putative target genes of unique super-enhancer domains in lymphoblast and frontal cortex

**Table 2.** GO analysis of putative target genes of common super-enhancer domains

| Term | Fold Enrichment | p-value |
| --- | --- | --- |
| GO:0098609~cell-cell adhesion | 4.14 | 4.69E-18 |
| GO:0000122~negative regulation of transcription from RNA polymerase II promoter | 2.56 | 2.64E-15 |
| GO:0045944~positive regulation of transcription from RNA polymerase II promoter | 2.14 | 8.07E-12 |
| GO:0045892~negative regulation of transcription, DNA-templated | 2.17 | 2.42E-05 |
| GO:0006351~transcription, DNA-templated | 1.52 | 2.19E-05 |
| GO:0006366~transcription from RNA polymerase II promoter | 2.01 | 5.22E-04 |
| GO:0071364~cellular response to epidermal growth factor stimulus | 6.57 | 4.60E-04 |
| GO:0030036~actin cytoskeleton organization | 3.20 | 0.0011 |
| GO:0008360~regulation of cell shape | 3.10 | 0.0010 |
| GO:0048870~cell motility | 7.53 | 0.0011 |
| GO:0008283~cell proliferation | 2.12 | 0.0020 |
| GO:0008285~negative regulation of cell proliferation | 2.05 | 0.0025 |
| GO:0045893~positive regulation of transcription, DNA-templated | 1.90 | 0.0027 |
| GO:0043547~positive regulation of GTPase activity | 1.82 | 0.0042 |
| GO:0043433~negative regulation of sequence-specific DNA binding transcription factor activity | 4.22 | 0.0046 |

### Genomic properties of common super-enhancer domains

To reason that super-enhancers play a key role in gene regulation, one would expect to see a strong correlation between the activity of a super-enhancer domain and the expression of its putative target gene. In general, putative target gene expression was significantly correlated with activities of super-enhancer domains (one-sample t-test, p-value < 10^−15^ for common, non-unique, and unique super-enhancer domains) (Figure 3a, see Methods). However, common super-enhancer domains showed the lowest correlation coefficients, when compared to unique and non-unique super-enhancer domains (KS test,*** p-value < 10^−15^). For example, a common super-enhancer domain shows considerable activity with enriched H3K27ac signals, but its putative target gene, MLL5, does not show a concordant expression level with the super-enhancer domain activity in all of the cell/tissue types (Figure 3b). This result raised the possibility that common super-enhancer domains are responsible for an additional biological role, other than the expression regulation of genes within their close proximity.

**Figure 3.**
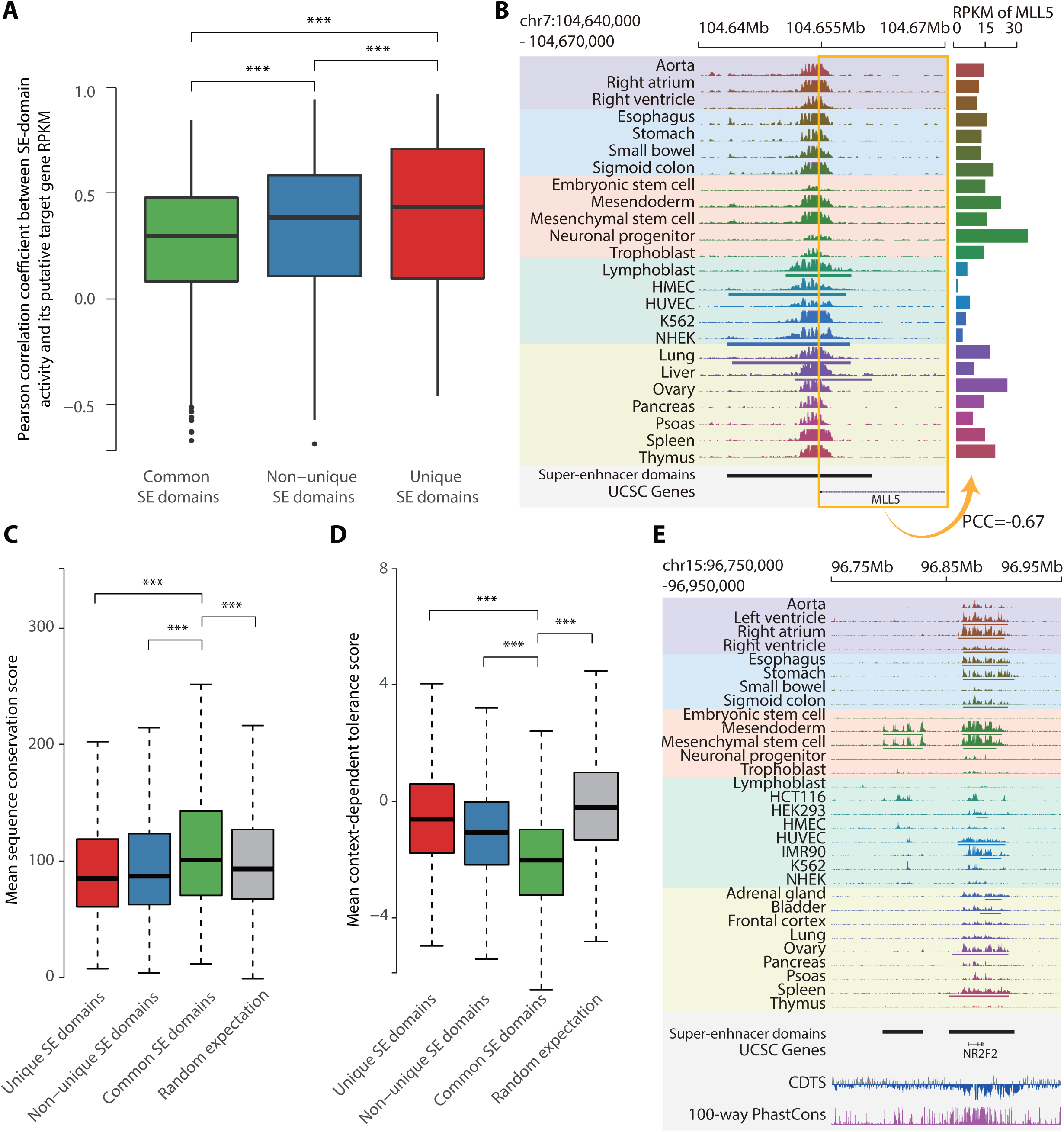
Genomic characterization of common super-enhancer domains. **a,** Boxplots of correlation between log2(H3K27ac RPM+1) in super-enhancer domains and RPKM of putative target genes (n=1,208 for common SE domains, n=3,733 for non-unique SE domains, n=1,098 for unique SE domains). Pearson correlation coefficient was calculated for each super-enhancer domain across 24 cell/tissue types. Significance of difference between super-enhancer domain categories was evaluated by KS test (*** p-value < 5×10^−12^). **b,** Example of common super-enhancer domain showing high H3K27ac signal across cell/tissue types but showing irregular RNA expression level. Left panel presents genome browser snapshot of H3K27ac signals at super-enhancer domain. Right panel presents RNA-seq RPKM of MLL5 gene. Pearson correlation coefficient (PCC) between H3K27ac signal at super-enhancer domain and MLL RPKM is shown below. **c,** Boxplots of mean sequence conservation score for each super-enhancer domain (n=1,098 for unique SE domains, n=3,733 for nonunique SE domains, n=1,208 for common SE domains). Significance of difference between super-enhancer domain categories was evaluated with KS test (*** p-value < 1.1 × 10^−5^ between random and others, p-value < 5.1 × 10^−12^ for the rest). **d,** Boxplots of mean CDTS for each super-enhancer domain (n=1,098 for unique SE domains, n=3,733 for non-unique SE domains, n=1,208 for common SE domains). Significance of difference between super-enhancer domain categories are evaluated with KS test (*** p-value < 1.7× 10^−9^ for unique and random, p-value < 3.0×10^−14^ for the rest). **e,** An example of adjacent non-unique super-enhancer domain and common super-enhancer domain illustrating evolutionary conservation of common super-enhancer domain. Shown together are 100-way PhastCons and CDTS. 100-way PhastCons track has the scale from 0 to 1, and CDTS track is scaled from −20 to +20. Common and unique super-enhancer domain are shown as right and left bar, respectively, in ‘Super-enhancer domains’ annotation.

Further investigation revealed that common super-enhancer domains are often located in evolutionarily conserved genomic regions compared to other super-enhancer domains and random expectation (Figure 3c, see Methods). However, we did not observe a strong conservation score in the unique super-enhancer domains compared to random expectation. Similarly, genomic areas of common super-enhancer domains also tend to have a low context-dependent tolerance score (CDTS)[9], which represents the frequency of sequence variation across the human population (Figure 3d, see Methods). The lower CDTS indicates a sequence is more resistant to the expected variation. Thus, low CDTS for common super-enhancer domains suggests, again, a potential function of these domains in universal genome regulation. For example, a common super-enhancer domain nearby NR2F2 gene presents both a high evolutionary conservation score and a low CDTS compared to the adjacent non-unique super-enhancer domain (Figure 3e).

### Striking association of common super-enhancer domains during early establishment of 3D chromatin loops

A previous study has revealed an association between super-enhancers and 3D chromatin structure[4], where 3D chromatin loops have a wide range of recovery rate following cohesin degradation. Loop domains with a fast recovery rate contained a greater number of super-enhancers compared to slow-recovering loops. We surmised that common super-enhancer domains play a significant role in the loop recovery since rapid recovery of 3D chromatin structure could be more critical in the universal genome regulation compared to the cell/tissue-type specific regulation.

To test our hypothesis, we examined the enrichment of 336 super-enhancer domains called from HCT-116 human colorectal cancer cell lines in both loop anchors (two contact loci) and loop domains (between two anchors). For each time point from 20 minutes to 180 minutes after the degradation of cohesin and subsequent start of recovery, we classified chromatin loops into fast, slow, and moderate-recovery loops by the intensity percentile (top 10% defined as fast recovery loop and bottom 10% as slow recovery loop) (Figure 4a upper panel, see Methods). The overlap between super-enhancer domain and loop anchor or loop domain was calculated as the measurement of super-enhancer domain enrichment (Figure 4a lower panel, see Methods). At an early time point (20 minutes after cohesin recovery), fast recovering loop anchors and domains were strikingly enriched in common super-enhancer domains but not in unique super-enhancer domains (Figure 4b upper panels). In contrast, slow-recovering loop anchors were significantly enriched in unique super-enhancer domains but not in common super-enhancer domains. In order to confirm the result not only for super-enhancer domains of HCT-116 but also of the 30 cell/tissue types, we further expanded the analysis and observed concordant results (Figure 4b, lower panels). This result supports the cell/tissue-type independent role of common super-enhancers in 3D chromatin loop formation.

**Figure 4.**
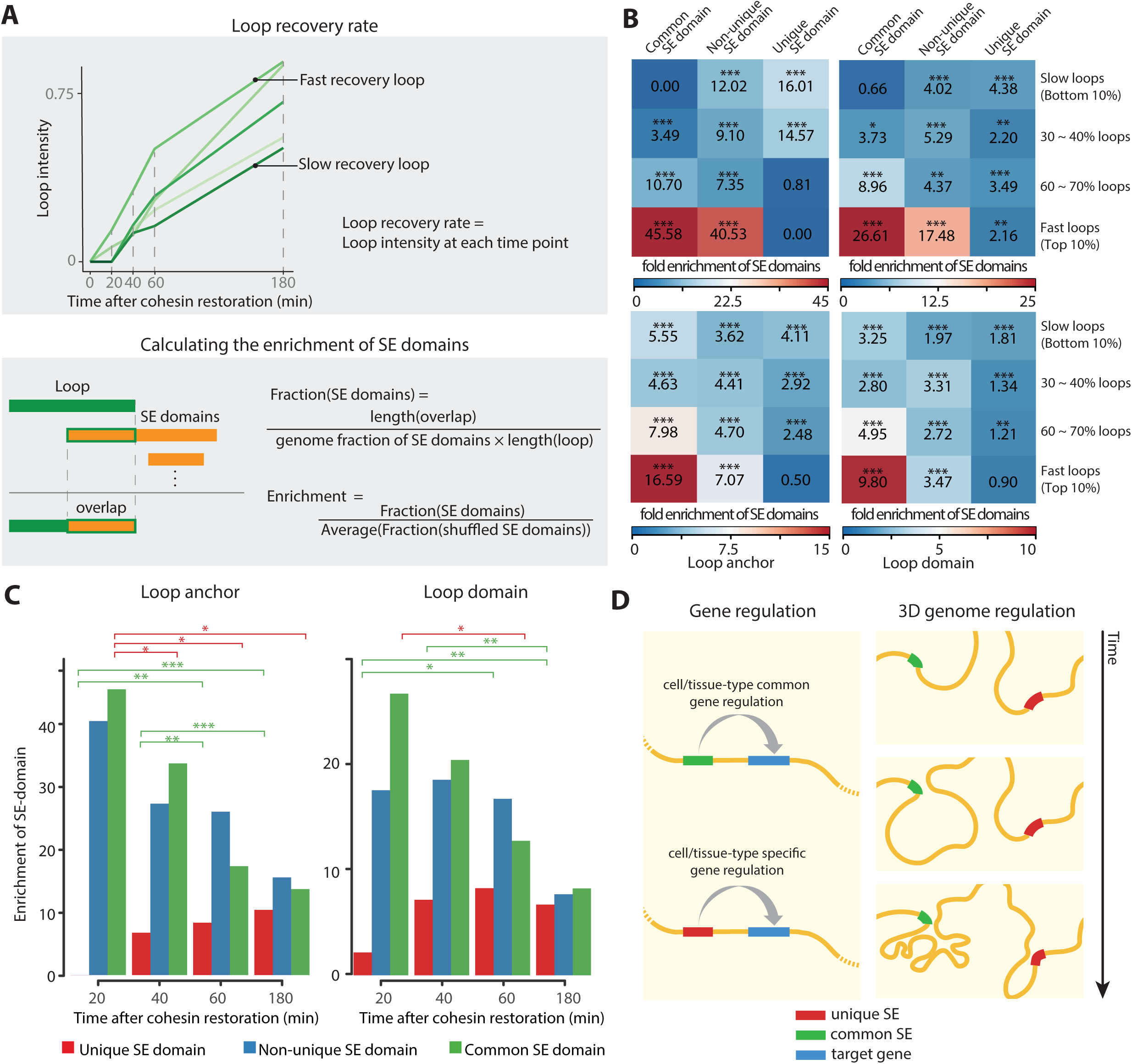
Common super-enhancer domains are associated with the early establishment of 3D chromatin loops. **a,** The schematic overview of calculated loop recovery rate (upper panel) and super-enhancer domain enrichment (lower panel). **b,** Heatmaps showing mean enrichment value of super-enhancer domains called from HCT-116 cell line (upper two panels) and all samples (lower two panels) at 20 minutes after cohesin recovery in loop anchors (left panels) and loop domains (right panels). Empirical p-values were calculated through random expectation values based on 10,000 iterations (* p-value < 0.05, ** p-value < 0.01, *** p-value < 0.001). **c,** Bar plots of super-enhancer domain enrichment in the top 10% fast recovery loops for each time point in loop anchors (left) and loop domains (right) (two-sample t-test, * p-value < 0.05, ** p-value < 0.01, *** p-value < 0.001). **d,** Model diagram of modes of super-enhancer functions. Canonical cell-type specific gene regulation (left) and 3D genome regulation of common super-enhancer domains (right).

We expanded the analysis to include later time points (see Methods), and revealed that an enrichment of common super-enhancer domains at both loop anchors and domains was the highest at the earliest time point (20 minutes), and gradually decreased in later time points (Figure 4c). On the other hand, enrichment of unique super-enhancer domains significantly increased in the later time points (two-sample t-test, * p-value < 0.05, ** p-value < 0.01, *** p-value < 0.001). Our result implies that common super-enhancer domains are often associated with the initial establishment of 3D chromatin loops compared to other super-enhancer domains.

## Discussion and Conclusion

We present a systematic approach to assess and characterize super-enhancers in a wide array of human cell/tissue types. The effective computation of background normalized super-enhancer activity, and the use of simple, yet powerful mathematical expression to evaluate the tissue specificity have led us to identify a new class of super-enhancers. These super-enhancer domains exhibit a high degree of universality across many cell/tissue types – a novel aspect of super-enhancers that has never been spotlighted in previous literatures. The universally active super-enhancers, which we defined as common super-enhancer domains, display a strong association with fast recovering chromatin loops after a sequential cohesin removal and restoration. Although the enrichment of super-enhancers at fast-recovering loops is previously reported[4], our analysis further reveals that common super-enhancer domains are strikinlgy enriched in the fast-recovering loops by more than 12-folds, compared to unique super-enhancer domains. The implications that our data bring forth to the function of super-enhancers in shaping chromatin organization may be outlined as follows. First, common super-enhancers may facilitate recruiting a substantial amount of structural proteins, such as cohesin and CTCF, to expedite loop recovery. A sequential model is also an interesting possibility, where cell-type nonspecific formation of fast recovering loops facilitated by common super-enhancer guide the genome folding in a stepwise manner to lead following cell-type specific loop formation. Hierarchical model where cell-type specific loop formation by unique super-enhancers requires the higher order loop formation promoted by common super-enhancers is also reasonable. Further experimentation will be required to test our hypothesis for the validation.

To conclude, we propose two distinct modes of super-enhancers; one is cell-type specific gene regulation mainly mediated by unique super-enhancer domains and second is 3D genome regulation mediated by common super-enhancer domains as a non-canonical function (Figure 4d). Although there is a limitation to generalize our conclusion since we investigated only one cell type, we shed light on common super-enhancers as a new potential mechanism underlying the early establishment of 3D chromatin loops.

## Methods

### Human cell/tissue types

In this study, 30 human cell/tissue types were examined. This includes H1 embryonic stem cell and its derived cell types (H1 embryonic stem cell, mesendoderm, mesenchymal stem cell, neuronal progenitor cell, and trophoblast), 8 cell lines (GM12878, HUVEC, IMR90, K562, NHEK, HMEC, HEK293, and HCT-116), and 17 tissue types (adrenal gland, aorta, bladder, dorsolateral prefrontal cortex, esophagus, gastric, lung, ovary, pancreas, psoas, right atrium, right ventricle, liver, sigmoid colon, small bowel, spleen, and thymus).

### Super-enhancer call

H3K27ac and input ChIP-seq reads were downloaded from Roadmap Epigenomics, ENCODE project and GEO[10, 11]. Sources are specified in Supplementary Table 3. Reads were aligned to the human reference genome (hg19 assembly) using BWA-mem[12]. Unmapped and poorly mapped (MAPQ < 10) reads were removed. PCR duplicates were also removed using Picard Markduplicates[13]. The remaining reads were then used to call peaks using MACS2[14]. Default parameters were used except q-value 0.01 was replaced with the p-value less than 1E-05. Super-enhancers were defined by stitching peaks using ROSE[2, 5] with default parameters, except TSS exclusion zone size was adjusted to 2,500bp. TSS exclusion was used because the H3K27ac signal is enriched for both active enhancers and promoters. Super-enhancers that span ENCODE blacklist regions were discarded.

### Calling super-enhancer domains

Super-enhancer domains were defined by merging the super-enhancers called for 30 samples using BEDTools (see Figure 1a). Super-enhancers with any overlap were merged into one super-enhancer domain.

### Cell/tissue-type specificity measurement with Tau value

Tau score was utilized to assess the cell/tissue-type specificity of super-enhancer domain activities and gene expression. Tau score is a measure commonly used to quantify tissue-specificity of expression as it enables a threshold-free evaluation of cell/tissue-type specificity. Tau score ranges from 0 to 1, where 0 indicates general and 1 indicates specific.Tau score is calculated as 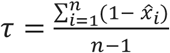; 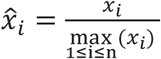[15], where x indicates cell/tissue-type specificity and n denotes the number of cell/tissue types.

### Gene expression data

Uniformly processed and quantified RPKM of protein-coding genes for 10 cell lines (H1 embryonic stem cell, mesendoderm, trophoblast, mesenchymal stem cell, neuronal progenitor cell, GM12878, HMEC, HUVEC, K562, and NHEK) and 14 tissues (aorta, esophagus, gastric, liver, lung, ovary, pancreas, psoas, right atrium and ventricle, sigmoid colon, small intestine, thymus, and spleen) was obtained from Roadmap epigenomics (“RNA-seq uniform processing and quantification for consolidated epigenomes”). The data had been normalized between cell/tissue types to have the same number of aligned reads on coding exons. Super-Enhancer domains called for none of the 24 cell/tissue types are excluded in gene expression analysis.

### Target gene assignment of super-enhancer domains

Putative target gene of a super-enhancer domain was defined as the gene with the closest transcriptional start site to the midpoint of the super-enhancer domain. For extended analysis, alternative definition utilized the 3 genes with transcriptional start site closest to the midpoint of super-enhancer domains.

### GO enrichment analysis using DAVID

For putative target genes of super-enhancer domains, GO terms were analyzed using DAVID 6.8[6, 16]. Significant GO_BP annotations were selected using p-value corrected by Benjamini-Hochberg procedure with threshold 0.05 and sorted by the p-value. For the analysis of the putative target genes of unique super-enhancer domains, we selected lymphoblast and frontal cortex as examples. Due to the small number of putative target genes of unique super-enhancer domains in each cell type, p-values tend to be insignificant. Thus, we presented up to top 10 enriched GO_BP annotations sorted with p-values without cutoff.

### Housekeeping genes

The putative target genes of super-enhancer domains were compared to the list of housekeeping genes provided by Eisenberg, E. *et al*[8] (http://www.tau.ac.il/~elieis/HKG/).

### Correlation between SE domain activity and expression level of its putative target gene

Pearson correlation coefficients between background-subtracted log2(RPM+1) value of H3K27ac ChIP-seq signal on super-enhancer domains and RPKM of their putative target gene were calculated across 24 cell/tissue. Super-enhancer domains called for none of the 24 cell/tissue types were excluded in the correlation analysis.

### Conservation and tolerance scores of super-enhancer domains

Evolutionary sequence conservation score calculated from multiple alignments of 100 vertebrates using phastCons method was obtained from the UCSC annotation database (phastCons100way). SumData column of 24bp resolution conservation score was mapped onto super-enhancer domains, and the mean value of mapped conservation score was used. CDTS referring to the variability of a nucleotide calculated for 11,257 human genomes was obtained from previous study[9]. CDTS of the 1bp resolution, was mapped onto super-enhancer domains and the mean of mapped CDTS was used.

### Hi-C matrix of HCT-116 following cohesin degradation and subsequent recovery

Hi-C matrix data of HCT-116 with 25kb resolution for control, cohesin degraded, and 20, 40, 60, and 180 minutes since subsequent recovery of cohesion, were obtained from the previous study (GEO accession number GSE104334)[4]. Each matrix was normalized by Knight-Ruiz algorithm. As the original read depth of each matrix was different, we normalized matrices to have the same total interaction frequency within 100kb distance since short-range interactions are known to be not affected by cohesin degradation. Chromatin loops were identified using HiCCUPS[17] with default parameters for medium resolution matrix. False positive chromatin loops from karyotypic abnormalities of HCT-116 cell line were filtered. The observed/expected enrichment filter of 4.5 was applied, as suggested by the previous study.

### Super-enhancer enrichment in loop anchors and domains

Recovery rate of loop anchors (interacting loci) and domains (region between two anchors) was defined as the ratio of loop strength at each time point compared to the strength in untreated cells (see Figure 4a, upper panel). Loops with monotonically increasing strength with regard to the time were selected for the analysis. Loop anchors were padded by 5kb in obtaining the super-enhancer domain enrichment. “Fraction” value of super-enhancer domains in loop anchors or domains was calculated by dividing the density of overlapping super-enhancer domains by the expected density (see Figure 4a, lower panel). The expected density of each class of super-enhancer domain is the proportion of the corresponding domains in the whole hg19 genome. The expected value of fraction for randomly shuffled super-enhancer domains is 1, with an infinite number of trials. The super-enhancer domain enrichment of a loop is obtained by dividing the observed fraction value by the random expectation of the fraction. The random expectation of fraction is the average of fractions calculated for 10,000 shuffled super-enhancer domains. Empirical p-value of a fraction where the distribution is defined as the result of 10,000 trials, is utilized as the significance measure of the enrichment. When comparing two enrichments, the p-value of two-sample t-test was utilized as the significance measure.

## List of abbreviations used

SE domain: super-enhancer domain
TSS: transcription start site
H3K27ac: histone 3 lysine 27 acetylation
ChIP-seq: chromatin immunoprecipitation sequencing
RPM: reads per million mapped reads
RPKM: reads per kilobase of transcript, per million mapped reads
CDTS: context-dependent tolerance score
KS test: Kolmogorov-Smirnov test

## Declarations

## Competing interests

None declared.

## Ethics approval and consent to participate

Not applicable.

## Consent for publication

Not applicable.

## Availability of data and materials

All data generated or analyzed in this study are included in this published article

## Funding

Ministry of Science, ICT, and Future Planning through the National Research Foundation in Republic of Korea [2017R1C1B2008838 to I. Jung]; Korean Ministry of Health and Welfare [HI17C0328 to I. Jung]. This work was supported by the KAIST Future Systems Healthcare Project from the Ministry of Science and ICT. Publication charges come from HI17C0328.

## Acknowledgments

We thank all members of Dr. Jung’s laboratory for their comments and criticisms on this study.

## Author contributions

JR, HK, and IJ conceived the study. JR led the data analysis with assistance from HK and DY. JR prepared the manuscript with assistance from AJL and IJ. All authors read and commented on the manuscript.

## Supplementary figure legends

**Supplementary Figure 1. Ubiquitous activity of super-enhancers defined in one cell/tissue type**

196 super-enhancer domains are called for only one cell/tissue type but showed constitutive H3K27ac signals across cell/tissue types and also located at less than 20% percentile of tau score. The heatmap of background subtracted log2 transformed H3K27ac signals of the domains across cell/tissue types is shown.

**Supplementary Figure 2. Hierarchical clustering of samples using H3K27ac signal on super-enhancer domains.**

Complete-link hierarchical clustering of distance between samples with respect to log2 transformed H3K27ac RPM of SE domains. H1 human embryonic cell line and its derived cell lines are shown in brown, and immortalized cell lines are shown in blue. Cancer cell lines are shown in bold text.

**Supplementary Figure 3. Functional characterization of super-enhancer domain classes with alternative putative target genes**

**a,** Heatmaps showing log2(RPKM+1) value of putative target gene of unique super-enhancer domains (top), non-unique super-enhancer domains (middle), and common super-enhancer domains (bottom). **b,** Histogram of tau score for putative target gene expression in each class of super-enhancer domains. Low tau score indicates universal expression pattern.

## Supplementary tables

**Supplementary Table 1.** GO analysis of alternative putative target genes of unique super-enhancer domains

|  | Term | Fold enrichment |
| --- | --- | --- |
| **GM12878**<br>**(Lymphoblast)** | GO:0006955~immune response | 3.6 |
|  | GO:0032729~positive regulation of interferon-gamma production | 11.3 |
|  | GO:0031295~T cell costimulation | 6.7 |
|  | GO:0033209~tumor necrosis factor-mediated signaling pathway | 5.0 |
|  | GO:0006915~apoptotic process | 2.3 |
|  | GO:0071222~cellular response to lipopolysaccharide | 4.6 |
|  | GO:0042832~defense response to protozoan | 13.7 |
|  | GO:0007165~signal transduction | 1.7 |
|  | GO:0060333~interferon-gamma-mediated signaling pathway | 5.5 |
|  | GO:0030154~cell differentiation | 2.3 |
|  | GO:0006954~inflammatory response | 2.4 |
|  | GO:0045893~positive regulation of transcription, DNA-templated | 2.1 |
|  | GO:0034138~toll-like receptor 3 signaling pathway | 24.4 |
|  | GO:0045630~positive regulation of T-helper 2 cell differentiation | 24.4 |
|  | GO:0040008~regulation of growth | 3.5 |
| **Prefrontal**<br>**Cortex** | GO:0007612~learning | 11.6 |
|  | GO:0010821~regulation of mitochondrion organization | 30.5 |
|  | GO:0008299~isoprenoid biosynthetic process | 28.3 |
|  | GO:0007409~axonogenesis | 6.7 |
|  | GO:0009240~isopentenyl diphosphate biosynthetic process | 132.2 |
|  | GO:0050992~dimethylallyl diphosphate biosynthetic process | 132.2 |
|  | GO:0060221~retinal rod cell differentiation | 88.1 |
|  | GO:0035434~copper ion transmembrane transport | 66.1 |
|  | GO:0006695~cholesterol biosynthetic process | 10.4 |
|  | GO:0060291~long-term synaptic potentiation | 10.4 |
|  | GO:0007010~cytoskeleton organization | 4.1 |
|  | GO:1901215~negative regulation of neuron death | 9.9 |
|  | GO:0055069~zinc ion homeostasis | 52.9 |
|  | GO:0071287~cellular response to manganese ion | 44.1 |
|  | GO:0009615~response to virus | 4.8 |

**Supplementary Table 2.** GO analysis of alternative putative target genes of common super-enhancer domains

| Term | Fold Enrichment | p-value |
| --- | --- | --- |
| GO:0098609~cell-cell adhesion | 2.353346006 | 1.92E-12 |
| GO:0000122~negative regulation of transcription from RNA polymerase II promoter | 1.620776655 | 1.18E-07 |
| GO:0045944~positive regulation of transcription from RNA polymerase II promoter | 1.480035009 | 1.92E-06 |
| GO:0008360~regulation of cell shape | 2.0838557 | 0.004124262 |
| GO:0007179~transforming growth factor beta receptor signaling pathway | 2.359877031 | 0.004662968 |
| GO:0030036~actin cytoskeleton organization | 2.035393939 | 0.015439556 |
| GO:0048008~platelet-derived growth factor receptor signaling pathway | 3.509299896 | 0.015539336 |
| GO:0001666~response to hypoxia | 1.853944092 | 0.02018621 |
| GO:0035556~intracellular signal transduction | 1.515181593 | 0.02630681 |
| GO:0048870~cell motility | 3.675016835 | 0.027716457 |
| GO:0045668~negative regulation of osteoblast differentiation | 2.957409997 | 0.031056171 |

**Supplementary Table 3.**
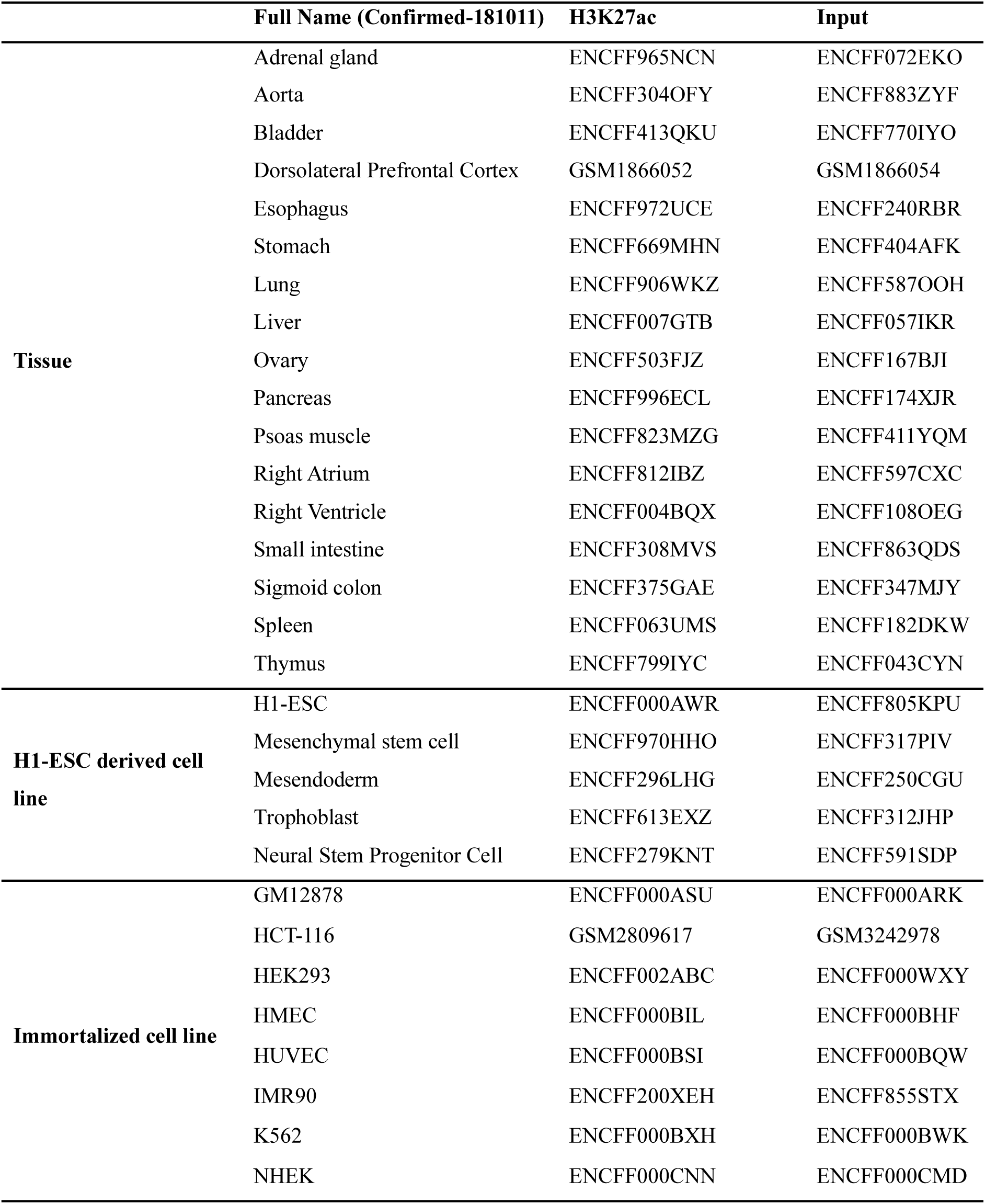
Source of H3K27ac and input ChIP-seq reads

